# Pathogen Detection in RNA-Seq Data with Pathonoia

**DOI:** 10.1101/2022.01.19.476681

**Authors:** Anna-Maria Liebhoff, Kevin Menden, Alena Laschtowitz, Andre Franke, Christoph Schramm, Stefan Bonn

## Abstract

**Motivation:** Recent evidence suggests that bacterial and viral infections may cause or exacerbate many human diseases. One method of choice to detect microbes in tissue is RNA sequencing. While the detection of specific microbes using RNA sequencing offers good sensitivity and specificity, untargeted approaches suffer from very high false positive rates and a lack of sensitivity for lowly abundant organisms.

**Results:** We introduce Pathonoia, an algorithm that detects viruses and bacteria in RNA sequencing data with high precision and recall. Pathonoia first applies an established k-mer based method for species identification and then aggregates this evidence over all reads in a sample. In addition, we provide an easy-to-use analysis framework that highlights potential microbe-host cell interactions by correlating the microbial to host gene expression. Pathonoia outperforms competing algorithms in microbial detection specificity, both on in silico and real datasets. Lastly, we present two case studies in human liver and brain in which microbial infection might exacerbate disease.

**Availability:** A Python package for Pathonoia sample analysis and a guided analysis Jupyter notebook for bulk RNAseq datasets are available on GitHub https://github.com/kepsi/Pathonoia.

**Graphical Abstract:** 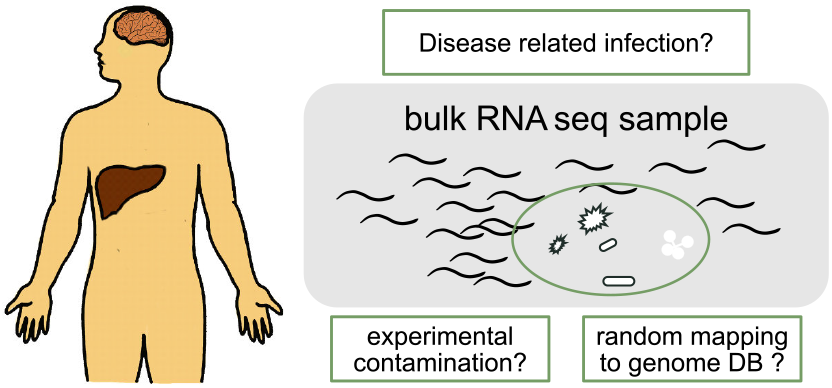

## 1 Introduction

A common approach to obtain insights into molecular mechanisms of disease is to sequence and analyze patient’s transcriptomes and compare them to transcriptomes of healthy controls. These experiments capture gene expression changes in human cells that might underly the disease process, but they can also capture transcripts of viruses or bacteria that infected those cells. While in many cases transcript information that cannot be aligned to the human genome or transcriptome is discarded as unspecific or contaminant, these non-human transcripts might be of microbial origin, and they might shed important disease insights.

Especially in recent years it has become clear that the human body harbors a vast amount of non-human cells. Certainly, these cells, mostly bacteria, are predominantly found in the gut and skin microbiomes, but other human tissues show abundance of microorganisms as well. For example, the healthy human blood microbiome is discussed by Castillo *et al*. (2019) and even the healthy brain is suspected to contain bacteria permanently according to Roberts *et al*. (2018).

For understanding the effect of certain agents in human tissues, dual RNA-sequencing experiments (Westermann *et al*. (2012)) are conducted *in vitro*. Nevertheless, it might be unknown that a pathogen relates to a disease. Real patient’s data is needed for discovering it and its co-morbidity, cause, or effects in the disease state.

The idea of finding foreign RNA in patient’s sequencing data has been proposed before, for example by Sangiovanni *et al*. (2019) and Rahman *et al*. (2018). The non-host part of a sample can be analyzed as a metagenome. Metagenomes, as known from the studies of microbiomes and environment, e.g. soil, come with their own challenges (Breitwieser *et al*. (2017)) but many tools exist to measure their abundance of bacteria and viruses, as proposed by Wood *et al*. (2019), Kim *et al*. (2016) and Alawi *et al*. (2019).

Many publicly available datasets have been analyzed with this notion by Simon *et al*. (2018) who created a database for a wide search of potential disease related pathogens. Similarly, we (Rahman *et al*. (2019)) created a database system for public small RNA experiments, that can be accessed online for a wide range of diseases and pathogens.

Nevertheless, the non-host part of a transcriptomic sample is very noisy, meaning that it contains majorly sequencing reads which do not have a biological background. These reads may instead stem from (human) processing contamination, bad sequencing quality and intentionally added sequences as part of the experimental protocol. The problem of falsely detected organisms (false positives) is a known problem for metagenomic data analysis (McIntyre *et al*. (2017)), but for the “metagenomes” which we observe here as a side-effect, this problem is oppressive.

Most metagenomic analysis tools measure pathogenic abundance by classifying for each sequencing read the organism it may stem from and summing up the reads according to their taxonomy. Analogous to the gene-count matrix, an organism count matrix is created. The problem with this approach is, that heavily (artificially) enriched sequences are often aligning to the same random organism and therefore provoking unreasonably high abundance. The traces of real pathogens may have a very low abundance and may drop out of the analysis. Furthermore, chimera, which are combined sequences of different species, cannot be (correctly) identified (Edgar *et al*. (2011)).

Therefore, we propose a solution which is considering the sample as a whole and measuring abundance of microorganisms across sequencing reads. Retaining an improved measure of abundance, common sequencing contamination can be differentiated from biological effects through the group-wise comparisons of their mean abundance.

## 2 Approach

Here, we are describing the Pathonoia algorithm, trying to overcome the problem of falsely detected organisms (false positives/FP) in a metagenomic sample. Our motivation originates in samples which contain a low number of true positives (TP) combined with many artificially contaminating and low-quality RNA sequences. For these samples it is especially hard to gain high specificity for the detected organisms.

Artificial sequences tend to result in many identical reads which are then being added up in read counts for a specific organism which they are randomly mapping to. In its evaluation step, Pathonoia excludes identical sequences from the counting since a higher abundance of a natural organism would result in a higher number of non-identical reads. Furthermore, longer matches to an organism’s genome are accounted with more weight than shorter ones.

Currently, Kraken’s metagenomic alignment is widely used and accepted as most efficient. Our proposal of Pathonoia is using Kraken’s k-mer matches to all existing bacterial and viral genomes in the NCBI database and replace its read classification step with a sample wide evaluation for organism abundance.

As displayed in Fig. 1F, with this approach Pathonoia is achieving greater precision on the detected organisms than Kraken’s read based evaluation alone and other Kraken-based abundance measuring techniques. We measured this precision on an artificial dataset since this is the only way to account for true false positives.

**Fig. 1.**
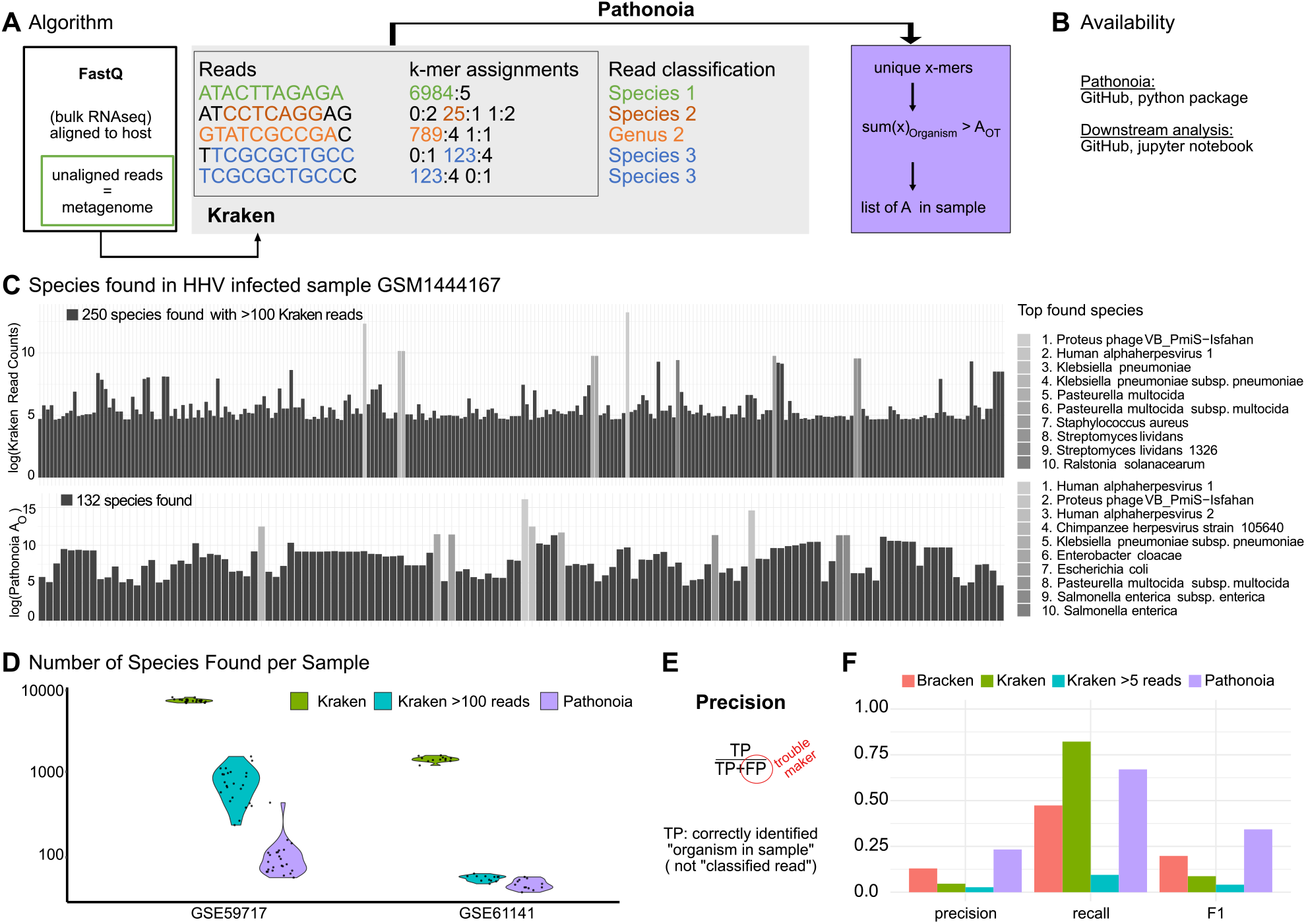
Pathonoia reduces number of false positives in noisy metagenomic samples. A) The Pathonoia algorithm is based on Kraken 2, analyzing unaligned reads in RNAseq data. Kraken generates k-mer assignments for each read, followed by a taxonomic classification (grey box). Pathonoia uses only the k-mer assignments of a whole sample and combines them into a non-read-count based abundance measure. Unique x-mers in the whole sample are identified, where an x-mer are several k-mers in a row, e.g., first read (green), 5 k-mers, *k* = 7, *x* = 11. For all unique x-mers belonging to the same organism, x is summed up into *A*_*O*_. If *A*_*O*_ *> A*_*OT*_, a threshold of approximately one full read-length or 3 distinct k-mers, the species is added to the list of organisms in the sample. B) Pathonoia and the downstream analysis template (Fig. 2) are available on GitHub. C) A spectrum of species reported in Kraken and Pathonoia, highlighting the top 10 most abundant species. Kraken reported 7262 organisms of which 250 organisms are shown, which have more than 100 reads for a cell line sample infected with Human Herpes Virus. Pathonoia reduces this list to 132 organisms and HHV and other Herpes viruses ascend in the ranking of reported species. D) Number of reported species with Kraken, Pathonoia and Kraken with threshold for two datasets with 12 and 24 samples. The threshold version counts an organism only as “detected” if it counts at least 100 reads. A lower number of detected organisms is desirable since that clears out the spectrum as shown in C) and reduces the number of FP. E) The aim of Pathonoia is to improve the precision of detected organisms to make better claims about the presence of unexpected organisms in a sample. FP (sequencing errors, other sample bias or random alignments, especially with poor quality reads) should be removed. Precision depends on TP and FP. True or false classification is defined by the presence of an organism in a sample. F) Based on a simulated dataset with 7 samples containing twelve to 50 species, precision, recall and F1 are measured for Kraken, Kraken with threshold, Pathonoia and Bracken. Kraken’s recall is the highest as it contains the highest number of species. Kraken is the base for all these algorithms and therefore they can only detect organisms which were detected by Kraken (with some exceptions, see Suppl. S1.1). With removing FP from the Kraken results, every algorithm also loses some TP, which is why recall goes down. Thresholding alone even worsens precision. For precision and the balanced F1 score, Pathonoia achieves best results. The downstream analysis (Fig. 2) helps to put samples and abundance measures in context of a dataset and can remove further sample bias.

The evaluation of a biological sample with Kraken-only and Pathonoia is visualized in Fig. 1C. Pathonoia is clearly able to reduce the number of (falsely) detected organisms, but also improves the profile of possibly abundant organisms, meaning the differences in “abundance counts” between included species.

Making use of Pathonoia’s abundance measures of species, we propose a downstream analysis for gaining biological results on a dataset (Fig. 1E). Here, we propose to compare case and control samples in a differential abundance analysis to uncover the potential involvement of a species in a condition.

## 3 Methods

First, we describe Pathonoia, which has the goal to uncover organic RNA-sequences from a mixture of artificial and metagenomic sequences. This set of “noisy” sequences may stem from the unaligned reads from RNA sequencing data from (human) transcriptome samples. Furthermore, our benchmarking methods are being explained.

Second, we describe an analysis pipeline based on the output of Pathonoia. Here, the goal is to distinguish biological signals from experimental and statistical contamination. We provide two exemplary studies with it. This pipeline is available as a guided analysis in a Jupyter Notebook on GitHub.

### Pathonoia

*Only non-host-mapping reads* are assumed as input for Pathonoia for detecting biological signal in noisy RNA-sequencing data. SAMtools can be used to assemble the corresponding fastQ files after using any aligner (see Suppl. Section S1 for further details). This input is processed in two major steps: metagenomic alignment with Kraken (Wood *et al*. (2019)) and Pathonoia’s sample-wide aggregation of k-mers.

*Identifying the lowest common ancestor (LCA) for each k-mer* in a sample is accomplished using Kraken. We use minimizer and k-mer lengths *l* = *k* = 31 for highest precision settings of Kraken 2 and the Kraken 2 index for all viral and bacterial genomes in the NCBI nt database. The LCA for a k-mer is the most specific taxonomic description of a set of organisms that share this sequence. We use the taxonomic identifier (taxID) for the implementation of the algorithm. The outcome of the alignment step is the *kraken-align* file, which contains the classified k-mers for each read of a sample. One example format of a classified read is:

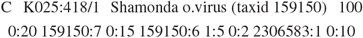

The five fields are:

1. *C* or *U* if the read was classified or not
2. the read identifier as given in the fastQ file
3. the taxonomic name and taxID with which it was classified according to the Kraken algorithm
4. the original read length
5. sequence of pairs in format taxID:Y, where Y is the number of k-mers in a row which are classified with the same taxonomic identifier

In this example, 20 k-mers could not be identified (taxID = 0), seven k-mers belong to the *Shamonda orthobunya virus* genome, followed by 15 unidentified k-mers and so on. Adding up the number of k-mers in a read, in this example 20 + 7 + 15 + 6 + 5 + 2 + 1 + 10 = 66, the result is always the read length subtracted by *k* + 1, here 100 − (31 + 1) = 66.

*Interpreting the Kraken alignment output* and detaching it from the notion of reads is the second and key step of Pathonoia. It is also described in Suppl. Fig. SF1. Using a hashmap, every identified sequence of a sample that is unique, is stored as key and its assigned taxID as value. We call these sequences (keys) as *x-mers*, since they have length *x* = *k* + *Y* (*k*: k-mer length, *Y* : number of consecutive k-mers identified with the same taxID). Next, all lengths *x* of sequences from the same organism (taxID) are summed up: 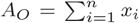, with *x* ∈ *X*, the set of *n* distinct x-mers of organism *O*. Only organisms surpassing a threshold of (our default) *A*_*OT*_ = 100 nucleotides (which corresponds to at least one fully mapping reads) are considered for the next step and final output. In order to increase certainty about a specific organism, the abundance measure *A*_*O*_ of every taxID on species level is summed with the abundance of genus and family levels additionally: *A*_*O*_ = *A*_*S*_ + *A*_*G*_ + *A*_*F*_. The intuition behind this is that if an organism is indeed part of the sample, several different areas of it are sampled and processed by chance. This may include areas which are not specific to the organism’s genome. If a species can be detected on species level though, the evidence can be increased by adding higher level counts which have to stem from a specific species in any case. Finally, the organisms can be ranked by their abundance *A*_*O*_. Nevertheless, the full potential unfolds with comparing sample groups with each other.

### Benchmarking Pathonoia

The aim of our algorithm is the reduction of falsely detected organisms in a metagenomic sample and therefore achieving as high precision as possible. Precision can be measured by evaluating a simulated dataset, where the presence (and absence) of every organism is known. Additionally, we evaluated a biological sample for testing Pathonoia qualitatively. We benchmark Pathonoia against read count-based abundance measuring techniques. We want to emphasize on comparing the technique of measuring abundance and do not intent to claim “improvements”, but rather adaptations for the aim of specific indication of lowly abundant organisms in a very noisy metagenomic sample.

*Read-count based abundance measures* are represented by Kraken 2 (Wood *et al*. (2019)) and Bracken (Lu *et al*. (2017)). Kraken 2 is the baseline for all algorithms in this benchmark. We ran all other abundance measure algorithms on top of its output, which makes the precision results comparable. No organism can be detected, if it wasn’t detected by Kraken 2 with at least one k-mer (i.e., there is a maximum value for TP and FP). Nevertheless, the way of counting and evaluating species differs in the various algorithms. Our benchmark includes “Kraken with cut-off minimum 5 reads”, which means that an organism counts as *detected* Kraken identifies minimum five reads originating from that same organism. The pure Kraken abundance counts an organism as *detected*, if at least one read is classified with it. Bracken is another tool based on Kraken which corrects abundance measures, including the statistical distribution of available genomes in the underlying database. Pathonoia may detect organisms with which no read was classified since it evaluates on k-mer level.

*For calculating precision, we used seven simulated samples*, which were constructed by Ye et al. (2019) for benchmarking taxonomy classifiers in metagenomics. They are samples containing DNA sequences from twelve to 50 organisms which are found in common human microbiomes or environmental metagenomes, for example the human gut or household. We oppose the common precision measurement techniques, which indicate if reads were correctly identified or not. Instead, we focus on the (in)correct presence of species. (Fig. 1E). Fig. 1F shows the average precision, recall and F1 score over all seven samples (detailed values in Suppl. Tables ST1-ST3). Overall, Pathonoia has an F1 score of 0.38 and the pure Kraken algorithm has 0.1, yielding an improvement of 0.34*/*0.09 ≈ 393%.

*Qualitative benchmarking* can be performed by looking at biological samples. The ground truth for the non-host sequencing reads is majorly unknown, which is why false positives and negatives cannot be determined. In Fig. 1C, We examined GEO sample *GSM1444167*, i.e. fibroblasts that were infected with *Human herpes virus* (HHV) *in vitro* Rutkowski *et al*. (2015). *In vitro* samples should be comparably clean, i.e., contain fewer contaminating organisms, but Kraken could still identify 250 species in the non-human reads which had over 100 reads. With Pathonoia, we could reduce this number to 132. Furthermore, the organism with highest abundance is the infecting Human alpha herpes virus 1 in the analysis with Pathonoia, where it was not the case for the Kraken analysis alone.

### Downstream Analysis

Applying Pathonoia on several samples in a dataset, results into a data matrix containing for each sample the abundance measures *A*_*O*_ of all organisms found in at least one of the samples. This abundance data, as well as some metadata about the samples, is the input for the downstream analysis. The goal is to identify differences of pathogenic abundance between sample groups, for example between diseased and control samples. This pipeline is realized as “guided analysis” and available online. In the following, we demonstrate the workflow exemplary on two datasets compare Fig. 2.

**Fig. 2.**
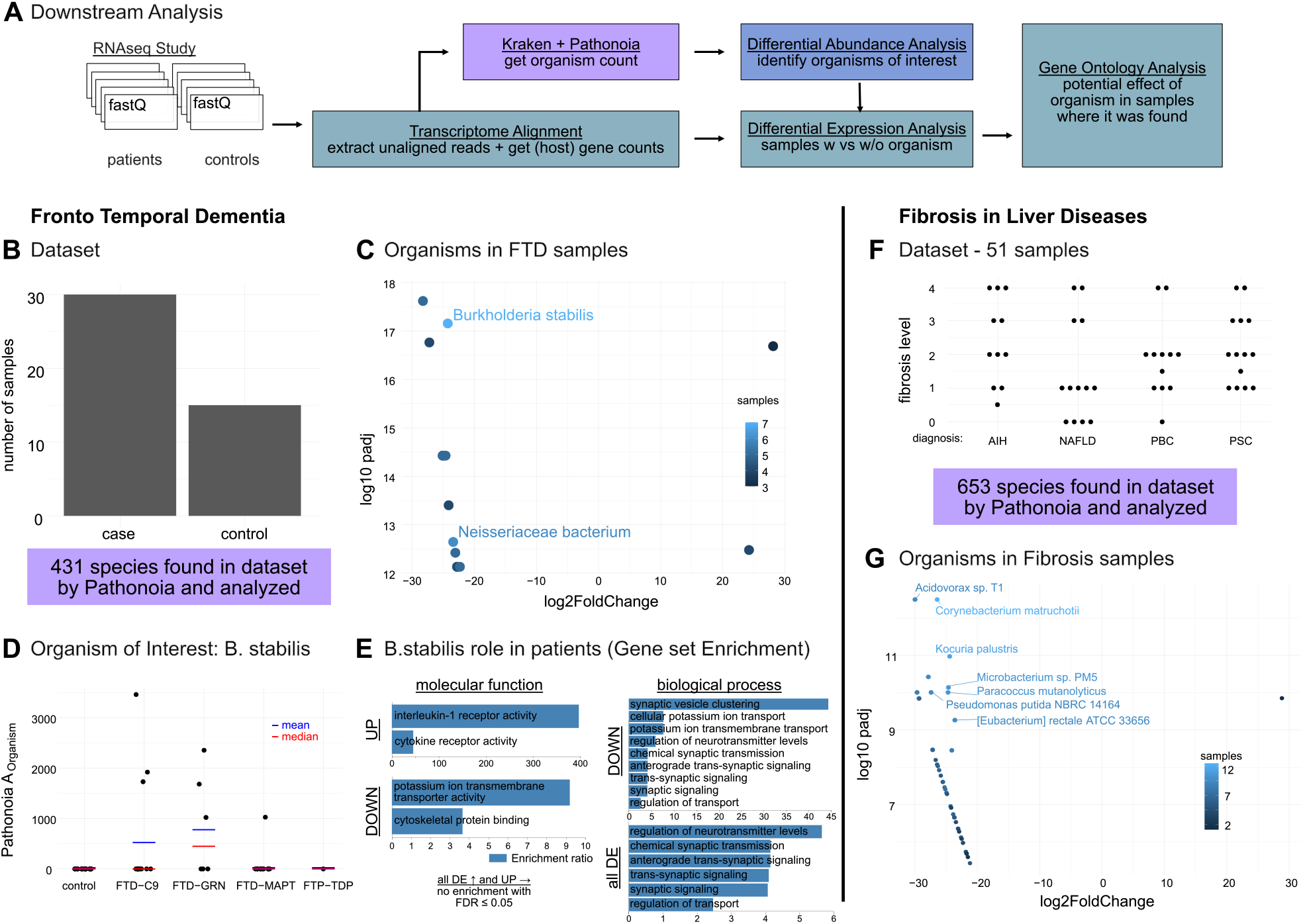
Analyzing datasets with Pathonoia. A) Analysis workflow for a dataset. Transcriptome alignment (input: fastQ files) yields gene counts and unaligned reads. Unaligned reads are analyzed by Kraken and Pathonoia as described in Fig. 1. A differential abundance (DA) analysis reports organisms that are more abundant in one sample group compared to another (examples in C, G). An “organism of interest” (OoI) can be selected for understanding its role in a sample group, e.g., patients. [Patient-] Samples with (*A*_OoI_ > 0) and without (*A*_OoI_ = 0) the OoI are selected for a differential gene expression (DE) analysis on the gene counts. Finally, a gene set enrichment analysis of up, down- or dis-regulated genes may uncover the pathways which the OoI affected. (B-E) Fronto Temporal Dementia B) The dataset contains 30 cases of FTD (sub-groups shown in D) and 15 controls. Pathonoia reported 431 organisms over all samples. C) The volcano plot shows 12 significantly (p-adj. < 0.05) DA organisms, ten of them up-regulated in FTD samples. The color scale shows the number of samples containing an organism. Only two organisms were found in more than 5 samples. D) Dots show *A*_*O*_ of B.stabilis per sample as it was present in most samples in C and therefore chosen as OoI. E) A DE analysis between FTD patients with and without B.stabilis lead to a gene set of 34 up-regulated and 109 down-regulated genes, in total 143. These three gene sets were compared with gene sets describing molecular functions and biological processes in a gene ontology analysis. The pathways’ enrichment ratios are shown with FDR≤ 0.5. [By B.stab] up-regulated genes in FTD patients hint towards an immune reaction in the patient and the biological processes relate to neural pathways. (F-G) Fibrosis in Liver Diseases F) A dataset with 51 human liver samples from patients with different liver diseases and fibrosis levels were analyzed with Pathonoia yielding 653 reported species over all samples. G) A DA analysis between non-fibrotic samples and samples with fibrosis level ≥ 1 lead to 41 significant (p-adj. < 0.05) DA organisms, where only one organism was up-regulated in two non-fibrotic samples. Seven organisms were present in more than nine fibrotic samples. A DE analysis between fibrotic samples with and without some of these organisms did not lead to any significant DE genes.

*The datasets* used for case study are comprising 48 and 63 samples. The first one is from an in-house study of Frontotemporal Dementia (FTD) (Menden *et al*. (2020)), looking at brain tissue samples split into disease and control according to Fig. 2B. The second dataset (Fig. 2F) contains liver samples from patients in different fibrotic liver stages due to one of four diseases: Autoimmune Hepatitis (AIH), Non-Alcoholic Fatty Liver Disease (NAFLD), Primary Biliary Cholangitis (PBC) and Primary Sclerosing Cholangitis (PSC).

*A Principal Component Analysis (PCA)* is executed as a first step of the analysis pipeline. It serves as an “outlier check”, where samples can be identified which may be overly contaminated. Furthermore, the PCA plot is colored by various available metadata (Suppl. Fig. SF3) for identifying if a certain experimental setting may result into major contamination. At this step, outlier samples are to be removed from the analysis (manually). In the FTD study we removed 3 samples and in the fibrosis study 12 samples were excluded. (Suppl. Fig. SF2) After re-executing the PCA for FTD and coloring according to age, gender, flow cell and RIN score, no bias could be pinpointed based on this metadata.

*The group-wise comparison* step includes the calculation of the mean abundance per organism for all samples in a group, for example all control samples. DESeq2 Love *et al*. (2014) is used for this differential abundance analysis. Originally, this tool was developed for differential gene expression analysis, but the same statistical model can be applied to our data. For increasing the stability of DESeq2, organisms which have zero abundance in most samples are excluded from the analysis. This is usually done for transcriptome analysis as well (Lin *et al*. (2016)). The output of this step is a list of organisms alongside their log2 fold change value between the sample groups and a p-value adjusted for multiple testing. A volcano plot colored by number of samples that an organism is present in, gives an overview of these results. (Results for FTD and fibrosis dataset in Fig. 2 C,G and Suppl. Tables ST4 and ST8)

*Abundance visualization* is the next step. The abundance is plotted per sample group for the most significant organisms. Here, it can be identified, if an organism plays a role in the whole sample group, or if the mean was only elevated due to one extreme case. The latter case should happen less often if more outliers were removed in the first step. In both studies, we observe that most organisms show consistent increase in the diseased samples but not in the controls (Suppl. Fig.s SF5 and SF7). The outcome of this step is the selection of an *organism of interest*. In the FTD study, we selected Burkholderia Stabilis, as it was the most prevalent organism of the differentially abundant ones. Its occurrence in the samples is visualized in the PCA plot in Suppl. Fig. SF4.

*A differential gene expression analysis* can help understanding if the organism of interest has a biological origin. Here, the primary transcriptome data is used for comparing samples that contain the organism and samples which do not contain the organism. These sample groups might be subgroup of another sample group. For instance, in the FTD dataset, we only select patient samples for this comparison, to understand which difference B.stabilis might make in the diseased case (and since it is not present in any control, also). Using DESeq2 again, on the original read counts, we retrieve a set of up- and down-regulated (DE-) genes. Ideally, this set concludes as an effect of the presence of the organism of interest. In the FTD study, this step resulted in a set of 143 significantly (p-adj. value < 0.05) DE genes. (More details in Suppl. Section S3)

*A Gene Ontology (GO) analysis* can be performed based on these gene set(s) for understanding the effect of the organism of interest. We perform the gene set enrichment analysis with WebGestalt Wang *et al*. (2017). (For the exact setting, see Suppl. Section S2). For the FTD study, all 34 upregulated genes were compared to the gene sets in the biological process database (Suppl. Table ST7). Most of the top ten enriched gene sets relate to pathways concerning the immune system and the results indicate an anti-bacterial response in brain tissue triggered by B.stabilis.

## 4 Discussion

*In an RNA-seq sample*, a fraction of sequencing reads does not align to the host’s transcriptome. Pathonoia was developed for making use of this situation and detecting organisms in this data. We try to find further information about the biological sample and evidence of potential infections in human tissue samples.

*Pathonoia’s precision* exceeds the one of methods which were adopted for this task previously. Those methods, not being designed for this task, measure abundance based on read counts. We compared our algorithm against other commonly used Kraken-based abundance measuring techniques and showed a reduction of false positives.

For all simulated samples, used for benchmarking, the recall is significantly higher than precision because of the high number of organisms found with Kraken. Most TP are discovered (high recall, 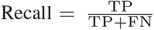, but also a high number of FP are detected, which in turn reduces precision (Fig. 1E). For improving certainty about Kraken-detected organisms, a cut-off can be introduced (“Kraken min five reads” in Fig. 1F) for reducing random hits. As a result, TP are reduced and both, recall and precision are dropping. Having an alternative look at Bracken, it does perform better than Kraken and is closer to Pathonoia’s values.

For the actual read-classification task, which Kraken and other tools were built for, precision and other performance measures are usually over 95% Ye *et al*. (2019). We showed, that with measuring performance based on the finding of specific organisms instead of correct classification of reads, the F1 score drops to less than 20% with common techniques. Pathonoia, in comparison, reaches up to 30%, which is much better but still not practical if used as stand-alone solution. We hope to start a new direction for the development of tools for gaining results with better biological interpretability. Pathonoia shows the best performance for the task of organism *detection*. With it we suggest a methodology, that is based on measuring abundance on sample level through adding up distinct sub-sequences of nucleotides for the aim of specific indication of abundant organisms in a very noisy metagenomic sample.

*Looking at real sequencing data*, we can observe various artifacts (Fig. 1C). Some organisms seem to be present in many samples, even though their presence was never mentioned or explained in the original study. They can be detected not only with Pathonoia, but also other algorithms. The most questionable one is *Proteus phage VB PmiS-Isfahan*. Phages are frequently used in biotechnical applications Kortright *et al*. (2020) and may serve for a quality check for the sequencing data. However, no evidence can be found for this hypothesis.

In an HHV-infected *in vitro* sample, Pathonoia detected three different herpes virus strains in the top hits, while Kraken could only detect the major one, *Human alpha herpes virus 1*. Furthermore, *Proteus phage VB PmiS-Isfahan* was the top hit for the Kraken algorithm. Its genome is relatively short (3.8 MB), and it was not mentioned to be part of the sample. *Klebsiella Pneumonaie* and *Pasteurella multocida* were found through both algorithms with relatively high abundance and are species who commonly appear in our environment Brown and Seidler (1973); Weber *et al*. (1984). They may have entered the sample during its collection, transport or manual processing. *Salmonella enterica* and *Staphylococcus aureus* on the other hand are extensively researched and sequenced organisms which inhabit and infect the human body (Knodler and Elfenbein (2019),Taylor and Unakal (2021)). Nonetheless, it cannot be determined which of these organisms was actually in the sample or contaminated it during the handling of a human being, for example in a hospital setting. Since this sample was infected *in vitro*, it usually implies, that it is comparably clean. Pathonoia can detect the known infection, but also helps to reduce the noise of random computational hits.

*For increasing certainty* about the detected organisms, samples should be considered in the reference of a whole dataset. Factors like sampling bias and sequencing noise may shadow the observations made on a single sample. We propose a downstream analysis comparing sample groups of a dataset to reduce the amount of sequencing artifacts and only find organisms which are unique to a defined condition. Furthermore, we suggest analyzing the transcriptome data of the host, from which our data was derived, in the light of the results of that comparison. Comparing the transcriptome of samples containing a bacterium versus samples not containing it within the same condition can give important indications on which effect the bacteria can have on the host.

Pathonoia and the downstream analysis are provided online on https://github.com/kepsi/Pathonoia. It produces several plots for an exploratory analysis. Some decisions must be taken manually, as for example the selection of an organism of interest once displayed with their differential abundance. Analyzing the results carefully is important, since a group difference in pathogenic load can come as well from a badly designed or executed experiment. Yet, to understand if the organism of interest has a biological origin, the transcriptome data can be used.

*An FTD case study* indicates that some patients may have (had) a *B*.*stabilis* infection in the brain. It was shown in literature that members of the *Burkholderia cepacia complex*, to which *B*.*stabilis* belongs, were able to infect human brain tissue and cause brain abscesses Hobson *et al*. (1995) and meningitis Peralta *et al*. (2018). Furthermore, *Burkholderia pseudomallei*, belonging to the same genus, are up-regulated in two independent, publicly available datasets concerning neurodegenerative diseases as shown in the Small RNA Expression Atlas Rahman *et al*. (2019) platform (Suppl. Table ST5). Nonetheless, it remains unclear whether *B*.*stabilis* plays an important role in FTD or could enter the brain for example due to already degenerated brain tissue. Outbreaks of *B*.*stabilis* in hospitals are reported frequently Heo *et al*. (2008); Martin *et al*. (2011). There was a major outbreak in Swiss hospitals due to contaminated washing gloves in 2016 Sommerstein *et al*. (2017). It is not known if patients from this cohort were treated there or in another hospital which had an unreported outbreak and if they could have been infected there. Further experiments should be conducted, such as dual RNA-seq experiments Westermann *et al*. (2012), which may help to answer, for example, the question how brain cells get affected by *B*.*stabilis*’ presence, and if it may be able to cause disease or change its progression.

*Fibrotic livers* are prone to bacterial infection through translocation from the gut (Ponziani *et al*. (2018)) and we could observe this in our samples as well. We found 41 species significantly differentially abundant where only one of them was highly abundant in 2 non-fibrotic samples. For the three most significant organisms, we conducted the human transcriptome differential expression analysis between fibrotic samples with and without the species, but it did not show any DE genes (for two species, Acidovorax sp.T1 and Kocuria palustris) and no enriched pathways for 19 DE genes of the third organism (Corynebacterium matruchotii). This may support the hypothesis, that liver fibrosis is not a reaction to infection, but rather allows infection to happen.

## 5 Conclusion

Our aim was to make use of the non-host part of RNA sequencing experiments and find potential infections or microbial abundance in the tissues under study. It is the nature of lowly abundant organisms that any algorithm cannot detect them with high certainty. Many random hits lead to noise in the data. With our proposed algorithm Pathonoia, it is possible to polarize some organisms from the noise. In contrast to the aim of other metagenomic algorithms, we focus on the detection of organisms, i.e., answering the question if an organism is present in the sample at all. Also, we wanted to overcome the commonly high false positive rate, which is especially increased in the kind of data we are focusing on. By viewing the full sample instead of individual sequencing reads, we reach almost 400% improvement in precision and uncovered pathogenic traces from noisy data.

We developed an algorithm for the analysis of non-host RNA sequencing reads which is specialized on detecting lowly abundant organisms. Furthermore, we proposed a downstream analysis for detecting microbiotic abundance in a group of samples within a dataset and for suggesting their influence on the host’s transcriptome. Two case studies give examples of the added value of our algorithm Pathonoia. They show that the developed algorithm can model biological context and may be able to support building new hypotheses and getting insights to disease.

## Acknowledgements

We thank Sven Heins, Sergio Oller and the ZMNH IT for their continuous technical support. We thank the members of the Institute for Medical Systems Biology for their helpful discussions and suggestions. We thank the team around Peter Heutink from the DZNE Tübingen for the scientific exchange and support around the FTD data.

## Funding

This work was supported by DFG CRU306 P-C to AL and LFF 78 to SB.

## Notes

https://github.com/kepsi/Pathonoia

### Competing Interest Statement

The authors have declared no competing interest.

