## Supplementary Material for "Pathogen Detection in RNA-Seq Data with Pathonoia"

### Supporting Information

#### S1 Pathonoia Algorithm Details

The Pathonoia algorithm takes FASTA or FASTQ files as input, which are evaluated by the Kraken 2 algorithm. It was developed for analyzing the adhering metagenome of an RNA-seq sample or dataset. For obtaining the unaligned reads from a host-aligned sample, SAMtools can be used on the BAM file produced by most aligners as follows:

```
samtools aligned.bam view -hb -f 4 > hostUnmapped.bam
samtools view hostUnmapped.bam | awk 'OFS="\t"; print ">" $1 "\n" $10' - > hostUnmapped.fa
```

The Kraken 2 index used for Pathonoia was built in March 2019 using the following commands:

```
./kraken2-build –download-taxonomy –db db/bacvir_k31
./kraken2-build –download-library bacteria –db db/bacvir_k31 –use-ftp
./kraken2-build –download-library viral –db db/bacvir_k31 –use-ftp
./kraken2-build –build –db db/bacvir_k31 –threads 8 –kmer-len 31 –minimizer-len 31
./kraken2-build –clean –db db/bacvir_k31
```

We set the k-mer length equal to the maximal Minimizer length in Kraken 2 $k = l = 31$. This forces Kraken to use no optimization (in terms of memory usage) at the cost of precision. You may refer to the original Kraken 2 publication (Wood *et al.* (2019)) for details. Since the relatively short $k = 31$ (default is $k = 35$) allows for too many random (incorrect) matches to the index, Pathonoia only considers sequences, where at least $z > 4$ k-mers in a row were classified identically (not mentioned in figure). We use $z > 4$ for matching the original k-mer default length of $k = 35$, where our index was built with $k = 31$.

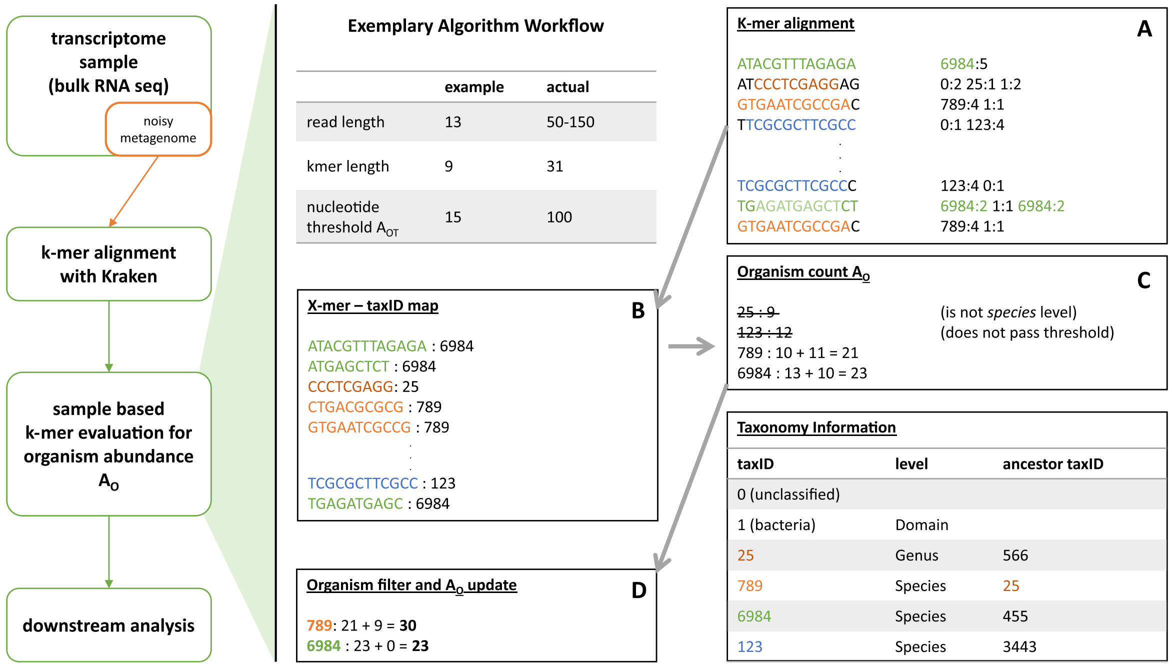


**Fig. SF1.** Pathonoia: Evaluation of all aligned k-mers in a sample. LEFT The non-host metagenome of a transcriptomic sample is k-mer-wise aligned to the NCBI nt database of all existing bacterial and viral genomes using Kraken. RIGHT (A) The aligned k-mers are (B) stored, connected as x-mers, in a hashmap with their corresponding taxID. (C) The lengths of x-mers with the same taxID are summed (D) The final abundance $A_O$ is calculated by adding genus and family summed lengths to the species summed length which passes a minimum threshold. (Figure originally published in Liebhoff (2021))

##### S1.1 Exceptions of discovered species, not discovered by Kraken

Kraken classifies each read with the Lowest Common Ancestor (LCA) of the species corresponding to all k-mers of this read. Pathonoia is based on the same species - k-mer alignments but abstracts from their read "environment". It counts the k-mers of the same species for all k-mers (in all reads) of the same sample. Therefore, it can theoretically be, that some species will not be reported by Kraken that is discovered by Pathonoia. In that case the LCA classification algorithm of Kraken is masking it.

#### S1.2 Recall, Precision and F1 Score for Benchmarked Algorithms

The following tables contain the underlying data for our benchmark and Figure 1F. They are recall, precision and F1 score for the seven simulated samples and their average per evaluated algorithm.

| sample | Kraken | K.>4 reads | Bracken | Pathonoia |
|---|---|---|---|---|
| buccal | 0.75 | 0.125 | 0.5 | 0.708 |
| cityparks | 0.929 | 0.082 | 0.48 | 0.673 |
| gut | 0.756 | 0.089 | 0.444 | 0.689 |
| hous1 | 0.785 | 0.092 | 0.446 | 0.661 |
| hous2 | 0.865 | 0.189 | 0.541 | 0.676 |
| nycsm | 0.739 | 0.065 | 0.413 | 0.609 |
| soil | 0.931 | 0.020 | 0.49 | 0.676 |
| mean | 0.822 | 0.095 | 0.473 | 0.670 |

**Table ST1.** Recall for simulated dataset comparing Pathonoia with Kraken algorithms

| sample | Kraken | K.>4 reads | Bracken | Pathonoia |
|---|---|---|---|---|
| buccal | 0.050 | 0.034 | 0.09 | 0.168 |
| cityparks | 0.053 | 0.030 | 0.137 | 0.274 |
| gut | 0.029 | 0.025 | 0.185 | 0.205 |
| hous1 | 0.049 | 0.038 | 0.176 | 0.285 |
| hous2 | 0.034 | 0.025 | 0.101 | 0.275 |
| nycsm | 0.073 | 0.030 | 0.157 | 0.226 |
| soil | 0.0376 | 0.004 | 0.062 | 0.197 |
| mean | 0.046 | 0.027 | 0.13 | 0.233 |

**Table ST2.** Precision for simulated dataset comparing Pathonoia with Kraken algorithms

| sample | Kraken | K.>4 reads | Bracken | Pathonoia |
|---|---|---|---|---|
| buccal | 0.093 | 0.059 | 0.152 | 0.272 |
| cityparks | 0.100 | 0.0430 | 0.214 | 0.389 |
| gut | 0.055 | 0.039 | 0.261 | 0.316 |
| hous1 | 0.092 | 0.054 | 0.252 | 0.398 |
| hous2 | 0.065 | 0.044 | 0.169 | 0.391 |
| nycsm | 0.134 | 0.041 | 0.228 | 0.329 |
| soil | 0.072 | 0.006 | 0.11 | 0.305 |
| mean | 0.087 | 0.041 | 0.198 | 0.343 |

**Table ST3.** F1 Score for simulated dataset comparing Pathonoia with Kraken algorithms

### S2 Downstream Analysis

For the analysis of our example datasets, we used the STAR aligner for retrieving aligned (and unaligned) reads using the following settings:

```
./STAR-2.5.3a/source/STAR –genomeDir ./indecies/STAR_Human/ –runThreadN 8 –alignIntronMax 0 –outFilterMismatchNoverLmax 0.06 –outSAMtype BAM SortedByCoordinate
–outStd BAM_SortedByCoordinate –outSAMunmapped Within –readFilesIn s1.fasta s2.fasta –outFileNamePrefix outdir/samplename_STAR_ > outdir/samplename_staralign.bam
```

In the final step of the downstream analysis, we use WebGestalt for the functional enrichment analysis of the (human) gene sets. We use the R version of the tool for the analysis of biological processes and molecular functions, with the following settings:

```
WebGes_BioProc <- function(genes, projectname)
    WebGestaltR(enrichMethod="ORA", organism="hsapiens", enrichDatabase="geneontology_Biological_Process", interestGene=genes, interestGeneType="ensembl_gene_id",
    referenceGeneType="genesymbol", referenceSet="genome_protein-coding", minNum=5, maxNum=2000, fdrMethod="BH", sigMethod="fdr", fdrThr=0.05,
    topThr=10, reportNum=20, perNum=1000, nThreads=64, isOutput=TRUE, outputDirectory="GO_results/GO_Analysis_biolProcess_FDR05",
    projectName=projectname, dagColor="continuous", hostName="http://www.webgestalt.org/")


WebGes_MolFun <- function(genes, projectname)
    WebGestaltR(enrichMethod="ORA", organism="hsapiens", enrichDatabase="geneontology_Molecular_Function", interestGene=genes,interestGeneType="ensembl_gene_id",
    referenceGeneType="genesymbol", referenceSet="genome_protein-coding", minNum=5, maxNum=2000, fdrMethod="BH",sigMethod="fdr",fdrThr=0.05,
    topThr=10, reportNum=20, perNum=1000, nThreads=64, isOutput=TRUE, outputDirectory="GO_results/GO_Analysis_molFunction_FDR05",
    projectName=projectname, dagColor="continuous",hostName="http://www.webgestalt.org/")
```

### S3 Supplementary Information for FTD study

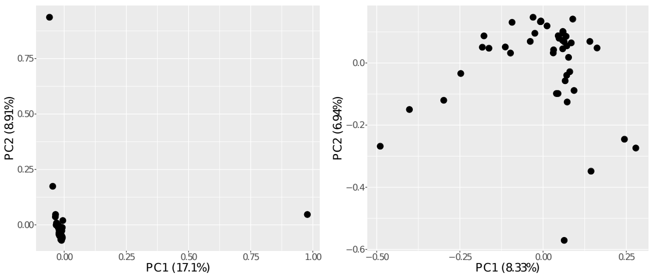


**Fig. SF2.** FTD PCA before (left) and after (right) 5 outlier removal.

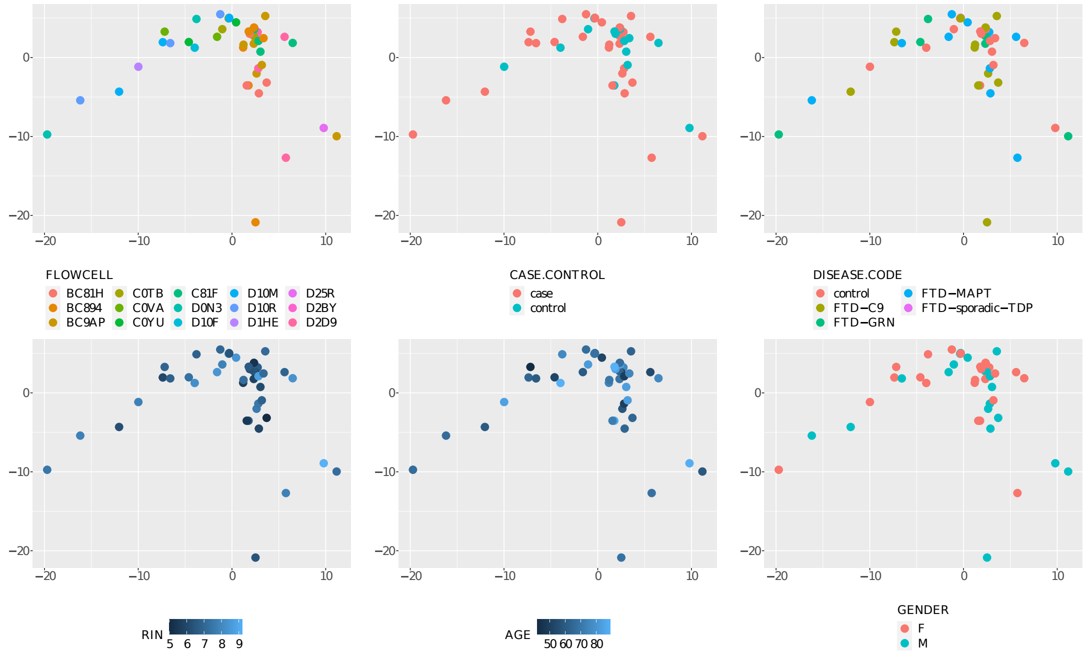


**Fig. SF3.** PCA of FTD dataset colored by different metadata. Clustering cannot be suggested based on the metadata.

| species name | taxID | base mean | log2FC | padj-value |
|---|---|---|---|---|
| Flavobacterium indicum | 1094466 | 957.01 | -28.88 | 1.7E-18 |
| **Burkholderia stabilis** | 95485 | 220.14 | -26.85 | 2.5E-18 |
| Caulobacter sp. FWC26 | 69665 | 454.84 | -27.85 | 1.2E-17 |
| Paracoccus yeei | 147645 | 152.09 | 27.49 | 1.5E-17 |
| Sphingomonas sp. PAMC26645 | 2565555 | 131.38 | -25.72 | 2.7E-15 |
| Burkholderia multivorans | 87883 | 181.25 | -25.69 | 2.7E-15 |
| Lactobacillus curvatus | 28038 | 49.67 | -24.75 | 2.9E-14 |
| Neisseriaceae bacterium | 2052837 | 27.47 | -24.02 | 1.6E-13 |
| Staphylococcus hominis | 1290 | 25.26 | -23.90 | 1.9E-13 |
| Arachidicoccus sp. KIS59-12 | 2341117 | 8.06 | 23.61 | 2.4E-13 |
| Acidovorax sp. 1608163 | 2478662 | 17.45 | -23.40 | 5.3E-13 |
| Rheinheimera sp. LHK132 | 2498451 | 17.20 | -23.38 | 5.3E-13 |

**Table ST4.** All significantly differentially expressed pathogens in brain tissue of FTD patients vs. control

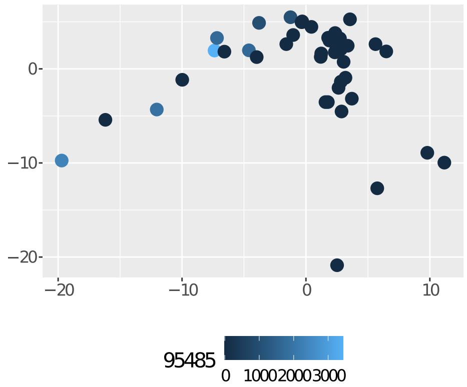


**Fig. SF4.** B. stabilis abundance coloring the PCA

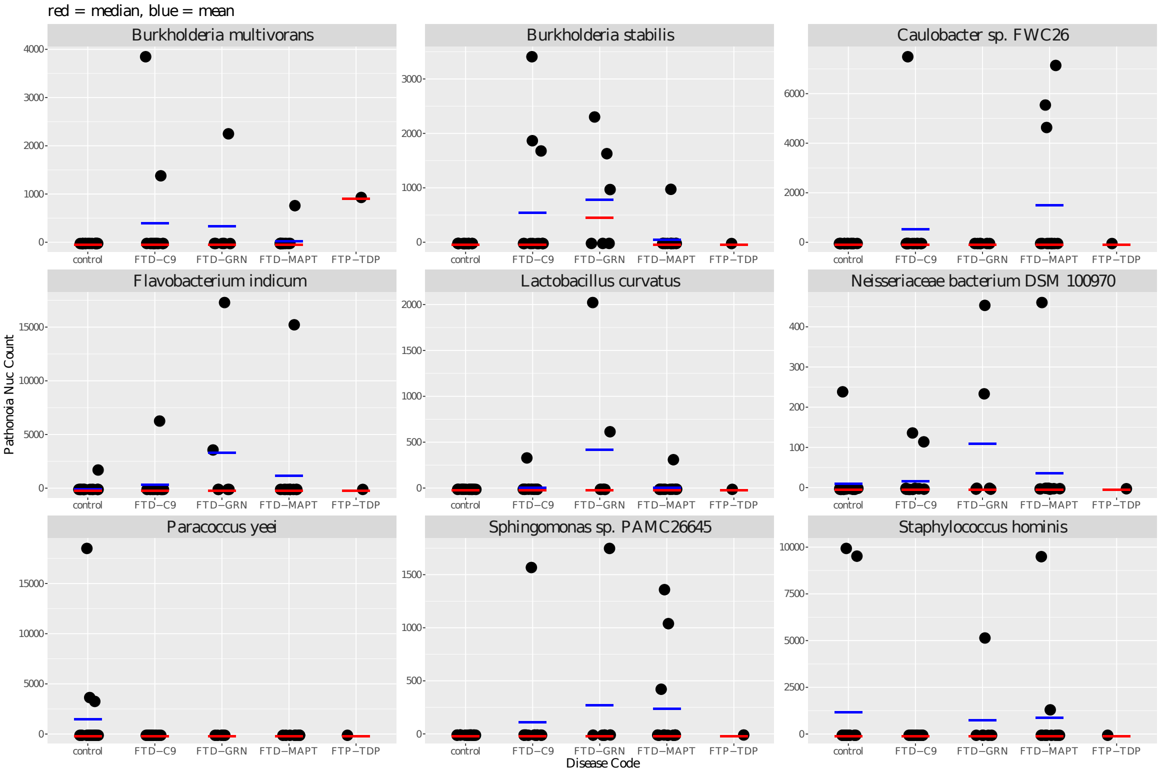


**Fig. SF5.** Top 9 differentially expressed organisms. The Pathonoia abundance for the nine most significant differentially expressed organisms between control and diseased samples is shown. Values are given for all disease phenotypes separately. Mostly the mean (blue bar) and median (red bar) abundance per sample is zero since presence of specific organisms is rare. Burkholderia stabilis is the most present organism in seven diseased samples. Even though the analysis was done comparing diseased and control samples, this plot shows the four different phenotypes of FTD. Here it can be seen how big values in single samples can lead to significant group differences. Nevertheless, some of the organisms show consistent increase in the diseased samples but not in the controls.

| Entity | Dataset | Condition | Group A | Group B | Covariates | log2 FoldChange | p-adj. value |
|---|---|---|---|---|---|---|---|
| Burkholderia pseudomallei | GSE46131 | disease | Lewy body dementia | non-demented | age-gender | -2.29 | 0.0048 |
| Burkholderia pseudomallei | GSE64977 | disease | healthy | Huntington's disease | age-gender | 1.68 | 0.0326 |

**Table ST5.** Using SEAweb with the search terms "brain" and "Burkholderia Pseudomallei" (closest to B.Stabilis as the latter was not found in SEA datasets), we find significant differential abundance of B. Pseudomallei in two independent datasets concerning neurological diseases comparing patient samples and control.

| Upregulated Genes | | | Downregulated Genes | | | | | | |
|---|---|---|---|---|---|---|---|---|---|
| MIR570 | CFAP47 | RGR | AC012184.2 | SYNPO | ZNF582-AS1 | BRSK1 | LSM10 | TBC1D25 | SKIL |
| CFAP45 | AL645608.1 | SNORA53 | ACRV1 | ELAVL3 | SEC16B | TCP11L1 | STX1B | DHX30 | ATP1A1 |
| AC018688.1 | AL596247.1 | MIR548H3 | SNORA50B | FAIM2 | RUNDC1 | KCNA2 | POGLUT1 | TTC7B | SCAMP5 |
| OTOG | GALNT4 | RN7SL32P | AC004706.4 | CDHR2 | GNB1 | PLEKHM3 | CD101 | VAMP2 | SALRNA1 |
| AQP4-AS1 | RN7SL525P | CD24 | MRGPRE | ACHE | LINC02192 | CAMK1D | MLST8 | RHOBTB2 | KCNN1 |
| HAUS4 | LINC01094 | PCAT5 | ERAL1 | NEFH | CLPTM1 | BTBD6 | HK1 | DMRTC1B | NOMO1 |
| GBP3 | CDC20B | RNF19A | EPB41L1 | ERICH1 | VWA7 | PCLO | AC068580.4 | PUM2 | PRRC2B |
| IL17RE | AC104058.1 | IL1RL1 | SYN1 | AES | AL009176.1 | KAZN | UBL7 | SLC29A1 | LARP1 |
| AL445483.1 | KIAA0391 | IL18R1 | PRDM8 | RNF157 | CISD3 | NOC2LP1 | NUDC | SEC16A | PRDM2 |
| RNU2-2P | P2RX4 | DOCK7 | HAPLN4 | TCEA2 | AAK1 | AL355075.4 | ZNF365 | SRP68 | HNRNPUL2 |
| AL513314.2 | SEC1P | | AC010970.1 | NCKIPSD | ZNF653 | AC244517.1 | SLC25A44 | ASPDH | KCNC3 |
| RF00017 | CBR3-AS1 | | PMS2P6 | KCNS1 | AC090587.2 | FP671120.3 | ZNF668 | CCNT2-AS1 | DYNLL2 |
| | | | CACNG8 | CUX2 | PSMC5 | MAEA | CYP1B1-AS1 | AL450992.2 | LZTS3 |
| | | | PI4K2A | TMEM39B | SLC9A1 | UBE2E2 | PHACTR1 | OPTN | GNG13 |
| | | | SSU72 | LRFN3 | CORO2B | BAAT | AKNAD1 | SIRPA | HAS1 |
| | | | GSK3A | NEFM | | | | | |

**Table ST6.** List of significantly (p-adj. value < 0.05) up- and down-regulated genes in FTD patient samples with vs. without B.stabilis infection.

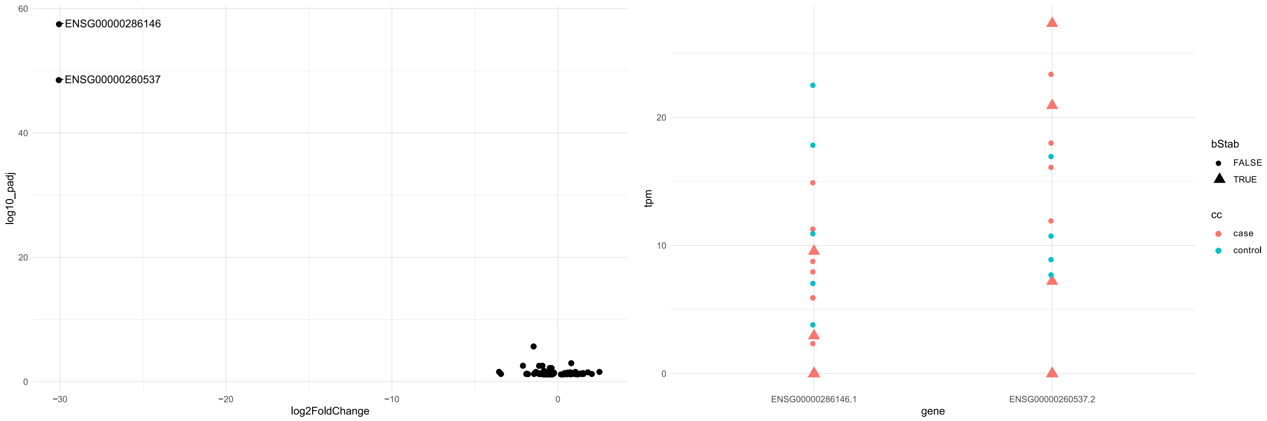


**Fig. SF6.** Diseased samples with and without B.stabilis were analyzed for differential gene expression, for understanding how it might be involved in the disease. LEFT 143 genes were discovered as significantly differentially expressed with an adjusted p-value lower than 0.05 (109 down regulated, 34 up regulated). Two down regulated genes show very high log2 fold change combined with an extremely low p-value. These genes are AC012184.2 (ENSG00000260537) and FP236383.9 (ENSG00000286146). RIGHT Their expression values (TPM) are given in the individual samples.
AC012184.2 is expressed in choroid plexus in the brain according to the Ensemble expression atlas Papatheodorou et al. (2019). According to its GeneCard Stelzer et al. (2016) it produces a novel, uncharacterized protein and is involved in ATP binding. For FP236383.9 much less information can be found. It was manually annotated by the Sanger Institute havana project Sanger Institute (2020), is uncharacterized and to be experimentally confirmed (TEC). According to MirTarBase Hsu et al. (2011) though, the miRNA hsa-mir-204-5p is targeting this gene. Using SEA Rahman et al. (2019), it can be found that this miRNA is associated with retinal degeneration Arora et al. (2010) and that the highest expression of this miRNA is found in eye and brain areas. Furthermore, various brain regions show up in the differential expression analysis results with high significance for this miRNA.

| geneSet | description | size | overlap | expect | enrichmentRatio | pValue | FDR |
|---|---|---|---|---|---|---|---|
| GO:0002825 | regulation of T-helper 1 type immune response | 27 | 2 | 0.021063 | 94.95157 | 1.95E-04 | 1 |
| GO:0042088 | T-helper 1 type immune response | 42 | 2 | 0.032765 | 61.04029 | 4.75E-04 | 1 |
| GO:0120163 | negative regulation of cold-induced thermogenesis | 47 | 2 | 0.036666 | 54.54664 | 5.95E-04 | 1 |
| GO:0071345 | cellular response to cytokine stimulus | 1015 | 5 | 0.791827 | 6.314513 | 7.08E-04 | 1 |
| GO:0034097 | response to cytokine | 1100 | 5 | 0.858137 | 5.826573 | 0.001022 | 1 |
| GO:0051241 | negative regulation of multicellular organismal process | 1143 | 5 | 0.891683 | 5.607376 | 0.001216 | 1 |
| GO:0019221 | cytokine-mediated signaling pathway | 705 | 4 | 0.549988 | 7.272886 | 0.001672 | 1 |
| GO:0032649 | regulation of interferon-gamma production | 95 | 2 | 0.074112 | 26.98623 | 0.002408 | 1 |
| GO:0032609 | interferon-gamma production | 106 | 2 | 0.082693 | 24.18578 | 0.002987 | 1 |
| GO:0032596 | protein transport into membrane raft | 5 | 1 | 0.003901 | 256.3692 | 0.003895 | 1 |

**Table ST7.** The top ten biological processes enriched by the set of 34 up-regulated genes in patients with B.stabilis present vs patients without this pathogen. The high FDR value for multiple test correction is less informative in this case, as the number of considered genes is low.

### S4 Supplementary Information for Fibrosis study

| tax ID | species name | log2FC | p-adj value | smpls |
|---|---|---|---|---|
| 1858609 | Acidovorax sp. T1 | -29.9827 | 2.93E-13 | 10 |
| 1114967 | Cutibacterium acnes TypeIA2 P.acn17 | -29.6547 | 9.04E-11 | 9 |
| 440085 | Methylorubrum extorquens CM4 | -29.3346 | 1.31E-10 | 7 |
| 28037 | Streptococcus mitis | -27.8456 | 3.48E-11 | 10 |
| 1211579 | Pseudomonas putida NBRC 14164 | -27.4035 | 9.04E-11 | 10 |
| 1636603 | Acinetobacter sp. ACNIH1 | -27.1565 | 3.14E-09 | 7 |
| 122355 | Pseudomonas psychrophila | -26.6323 | 5.95E-09 | 6 |
| 43768 | Corynebacterium matruchotii | -26.4355 | 2.93E-13 | 13 |
| 47885 | Pseudomonas oryzihabitans | -26.3994 | 7.87E-09 | 6 |
| 1879049 | Acinetobacter sp. WCHAc010034 | -26.2889 | 8.69E-09 | 7 |
| 401472 | Corynebacterium ureicelerivorans | -26.0778 | 1.12E-08 | 7 |
| 739141 | Methylobacterium sp. XJLW | -25.8098 | 1.56E-08 | 6 |
| 2219696 | Sphingomonas sp. FARSPH | -25.4983 | 2.31E-08 | 6 |
| 1325095 | Bradyrhizobium guangzhouense | -25.4678 | 2.31E-08 | 7 |
| 436515 | Variovorax boronicumulans | -25.2041 | 3.21E-08 | 8 |
| 2480908 | [Mycobac.] chelonae subsp. gwanakae | -25.0827 | 3.64E-08 | 8 |
| 47770 | Lactobacillus crispatus | -24.8951 | 4.54E-08 | 7 |
| 120107 | Sphingobium cloacae | -24.816 | 4.86E-08 | 6 |
| 1499308 | Paracoccus mutanolyticus | -24.6895 | 9.04E-11 | 11 |
| 2014534 | Microbacterium sp. PM5 | -24.6228 | 6.66E-11 | 11 |
| 71999 | Kocuria palustris | -24.4215 | 9.79E-12 | 12 |
| 2082188 | Sphingobium sp. YG1 | -24.2455 | 1.03E-07 | 7 |
| 1196325 | Pseudomonas putida DOT-T1E | -24.1474 | 1.13E-07 | 4 |
| 2282475 | Achromobacter sp. B7 | -24.0824 | 3.23E-09 | 9 |
| 162426 | Microbacterium hominis | -23.8223 | 1.70E-07 | 7 |
| 33010 | Cutibacterium avidum | -23.6556 | 2.05E-07 | 6 |
| 945844 | Massilia oculi | -23.6202 | 2.07E-07 | 6 |
| 515619 | [Eubacterium] rectale ATCC 33656 | -23.5969 | 5.00E-10 | 10 |
| 2079596 | Acinetobacter sp. SWBY1 | -23.3826 | 2.75E-07 | 5 |
| 29466 | Veillonella parvula | -23.055 | 4.10E-07 | 8 |
| 216778 | Stenotrophomonas rhizophila | -23.0221 | 4.15E-07 | 5 |
| 237610 | Pseudomonas psychrotolerans | -22.8647 | 4.94E-07 | 4 |
| 33033 | Parvimonas micra | -22.5676 | 7.04E-07 | 7 |
| 1945662 | Paracoccus contaminans | -22.4732 | 7.72E-07 | 4 |
| 1123269 | Sphingomonas sanxanigenens ... | -22.2548 | 9.91E-07 | 5 |
| 314722 | Pseudoxanthomonas suwonensis | -21.7936 | 1.72E-06 | 5 |
| 28131 | Prevotella intermedia | -21.676 | 1.92E-06 | 6 |
| 1177574 | Prevotella jejuni | -21.6626 | 1.92E-06 | 3 |
| 178339 | Actinomyces hongkongensis | -21.468 | 2.38E-06 | 3 |
| 1930557 | Shewanella sp. FDAARGOS_354 | -21.1338 | 3.48E-06 | 6 |
| 1804984 | Burkholderia sp. OLGA172 | 28.86658 | 1.31E-10 | 2 |

**Table ST8.** All significantly (p-adj < 0.05) differentially expressed pathogens in liver tissue of fibrosis patients vs. control (fibrosis level <1)

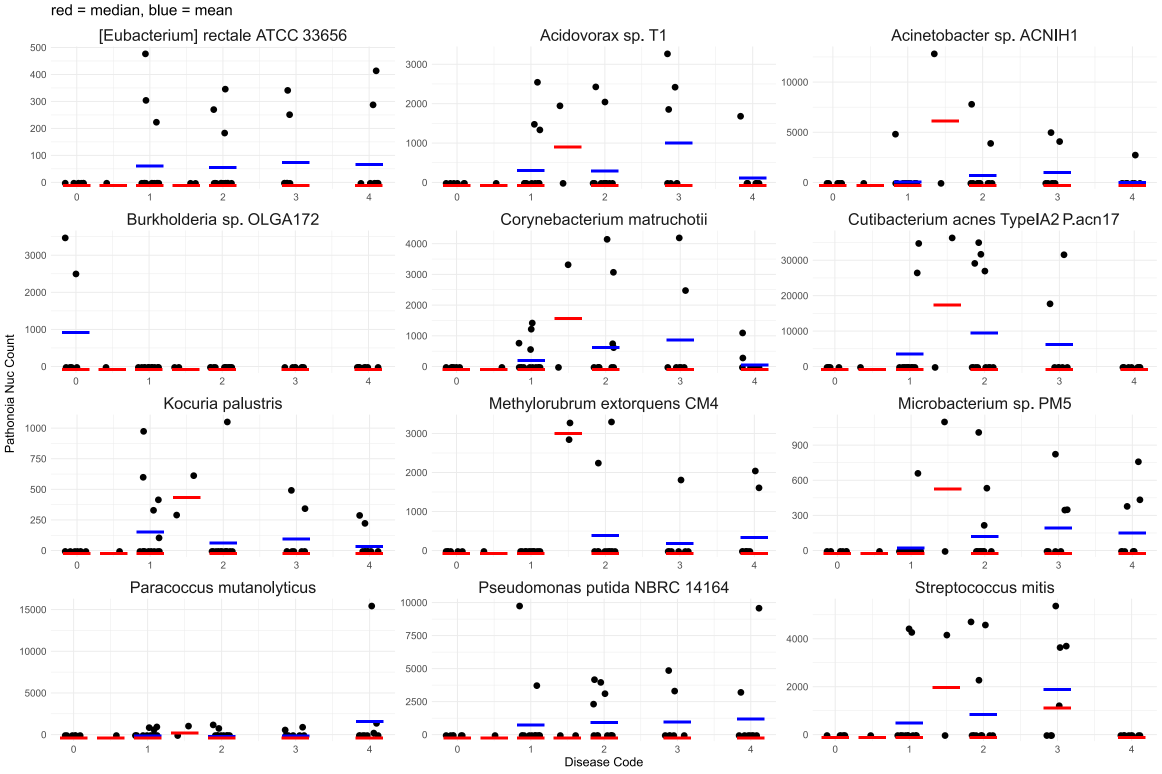


**Fig. SF7.** Top 12 differentially expressed organisms. The Pathonoia abundance for the nine most significant differentially expressed organisms between fibrosis and no fibrosis samples is shown. Values are given for all fibrosis levels separately. Mostly the median (red bar) abundance per sample is zero since presence of specific organisms is rare. Even though the analysis was done comparing fibrosis vs no-fibrosis samples, this plot shows all fibrosis levels. Only Burkholderia sp. OLGA172 shows abundance in non-fibrotic samples, but only two samples contain this pathogen.
